# Axon initial segment GABA inhibits action potential generation throughout periadolescent development

**DOI:** 10.1101/2023.04.16.537084

**Authors:** Anna M Lipkin, Kevin J Bender

## Abstract

Neurons are remarkably polarized structures: dendrites spread and branch to receive synaptic inputs while a single axon extends and transmits action potentials to downstream targets. Neuronal polarity is maintained by the axon initial segment (AIS), a region between the soma and axon proper that is also the site of action potential (AP) generation. This polarization between dendrites and axons extends to inhibitory neurotransmission. In adulthood, the neurotransmitter GABA hyperpolarizes dendrites but instead depolarizes axons. These differences in function collide at the AIS. Multiple studies have shown that GABAergic signaling in this region can share properties of either the mature axon or mature dendrite, and that these properties evolve over a protracted period encompassing periadolescent development. Here, we explored how developmental changes in GABAergic signaling affect AP initiation. We show that GABA at the axon initial segment inhibits action potential initiation in Layer 2/3 pyramidal neurons in prefrontal cortex from mice of either sex across GABA reversal potentials observed in periadolescence. These actions occur largely through current shunts generated by GABAA receptors and changes in voltage-gated channel properties that affected the number of channels that could be recruited for AP electrogenesis. These results suggest that GABAergic neurons targeting the axon initial segment provide an inhibitory “veto” across the range of GABA polarity observed in normal adolescent development, regardless of GABAergic synapse reversal potential.

## INTRODUCTION

Gamma-aminobutyric acid (GABA) is the primary inhibitory neurotransmitter in the neocortex. GABA binds to postsynaptic GABA_A_ receptors—ionotropic receptors that allow chloride and bicarbonate ions to flux across the membrane. The direction of current flow following GABA_A_ receptor opening is dependent on the concentration of chloride ions inside and outside the neuron. This distribution of chloride is regulated in part by cation-chloride cotransporters, which are dynamically expressed and regulated throughout development across neurons as well as across different compartments in the same neuron (Ben-Ari, 2002) (Báldi et al., 2010). Early in postnatal life, expression of the chloride importer NKCC1 is elevated, leading to high intracellular chloride levels and a depolarized GABA reversal potential (EGABA). In dendrites, NKCC1 is supplanted by KCC2, a chloride exporter, by the second postnatal week of life in mice (Ben-Ari, 2002). However, KCC2 appears to be largely excluded from axons in adulthood (Báldi et al., 2010; Hedstrom et al., 2008; Szabadics et al., 2006). This creates a gradient of chloride concentration and GABA reversal potential across the somatodendritic-axonal axis, ranging from a hyperpolarized E_GABA_ in dendrites to a more depolarized E_GABA_ in the axon.

The axon initial segment (AIS), the region of axon most proximal to the soma, is an intermediate zone separating somatodendritic and axonal domains. The AIS is the site of action potential (AP) generation under most conditions (Kole et al., 2008; Stuart et al., 1997). In pyramidal cells, the AIS is exclusively and selectively targeted by GABAergic synapses from chandelier cells (Somogyi et al., 1982; Veres et al., 2014). Based on this specific synaptic target, chandelier cells have been proposed to orchestrate synchronous activity across a network of pyramidal neurons by providing a powerful “veto” over pyramidal cell firing (Lewis, 2011; Veres et al., 2014). However, the functional effects of these synapses over development are not fully understood. In mature neurons (>4 weeks of age in mouse), GABAergic synapses at the AIS inhibit AP initiation, as E_GABA_is at a potential far more hyperpolarized than AP threshold (−85 – −90 mV) (Dugladze et al., 2012; Glickfield et al., 2009; Veres et al., 2014). But earlier in development, EGABA is more depolarized, mimicking levels more commonly observed in the distal axon in the first two weeks of life (Price and Trussell, 2006; Ruiz et al., 2010; Turecek and Trussell, 2001; Zorrilla de San Martin et al., 2017), before slowly hyperpolarizing over a periadolescent period (Rinetti-Vargas et al., 2017). Thus, chandelier cells are positioned to provide strong inhibitory control over action potential initiation in adulthood, but may instead have an excitatory role in early adolescence.

The postsynaptic effects of GABA are heavily dependent on E_GABA_. However, differences in EGABA alone do not fully determine whether a particular synapse has an excitatory or inhibitory effect. The effects of GABA are deeply context-dependent, relying on factors like resting membrane potential (V_m_), the position and timing of the GABA synapse relative to excitatory inputs, and intrinsic membrane conductances of the postsynaptic neuron (Gulledge and Stuart, 2003; Kwag and Paulsen, 2009; Lombardi et al., 2021). AIS GABAergic signaling provides a practical context to interrogate the effect of changing E_GABA_: these inputs display a change in EGABA at a point when other maturational changes are complete (Pan-Vazquez et al., 2020), they have a well-characterized postsynaptic target down to the subcellular level (Somogyi et al., 1982), and by virtue of their specific targeting, they provide an accessible output metric of inhibition: action potential initiation.

Here, we used imaging and electrophysiological approaches to characterize the consequences of AIS GABAergic input on AP properties across developmental E_GABA_ values. We found that GABAergic input to the initial segment, but not proximal dendrite, exerted an inhibitory effect on AP generation across E_GABA_ conditions. This inhibitory effect resulted from shunting inhibition and changes in voltage-gated sodium channel (Na_V_) availability for AP initiation, resulting in depolarization of AP threshold and delays in AP onset. Thus, GABAergic signaling onto prefrontal pyramidal cell initial segments is uniformly inhibitory across adolescent development, despite undergoing a developmental shift in E_GABA_ across this time period.

## RESULTS

### Axonal GABAergic input suppresses AP initiation at physiologically relevant E_GABA_ values

To understand how GABAergic inputs influence AP generation, we performed whole-cell *ex vivo* electrophysiology combined with GABA iontophoresis to probe AP probability in the absence and presence of GABA in L2/3 pyramidal neurons in prefrontal cortex slices from mice aged P24–P50. Intracellular chloride concentration was altered to set E_GABA_ within recordings to values measured in early adolescence (−50 mV) and adulthood (−90 mV), as determined in gramicidin-based perforated-patch recordings across development (**Fig. 1*A***) (Rinetti-Vargas et al., 2017). Neurons were filled via whole-cell dialysis with either the E_GABA_ = -90 mV or the E_GABA_ = −50 mV internal solution, both containing the dye Alexa Fluor 594 to allow for morphological identification of the AIS. The AIS was identified by its placement opposite the apical dendrite and its lack of spines. To bypass we delivered GABA via an iontophoretic pipette placed 20-30 μm from the soma, which corresponds to the site of AP initiation in these neurons (**Fig. 1*B***). Cells with axons isolated from neighboring basal dendrites were selected for subsequent experiments.

**Figure 1:**
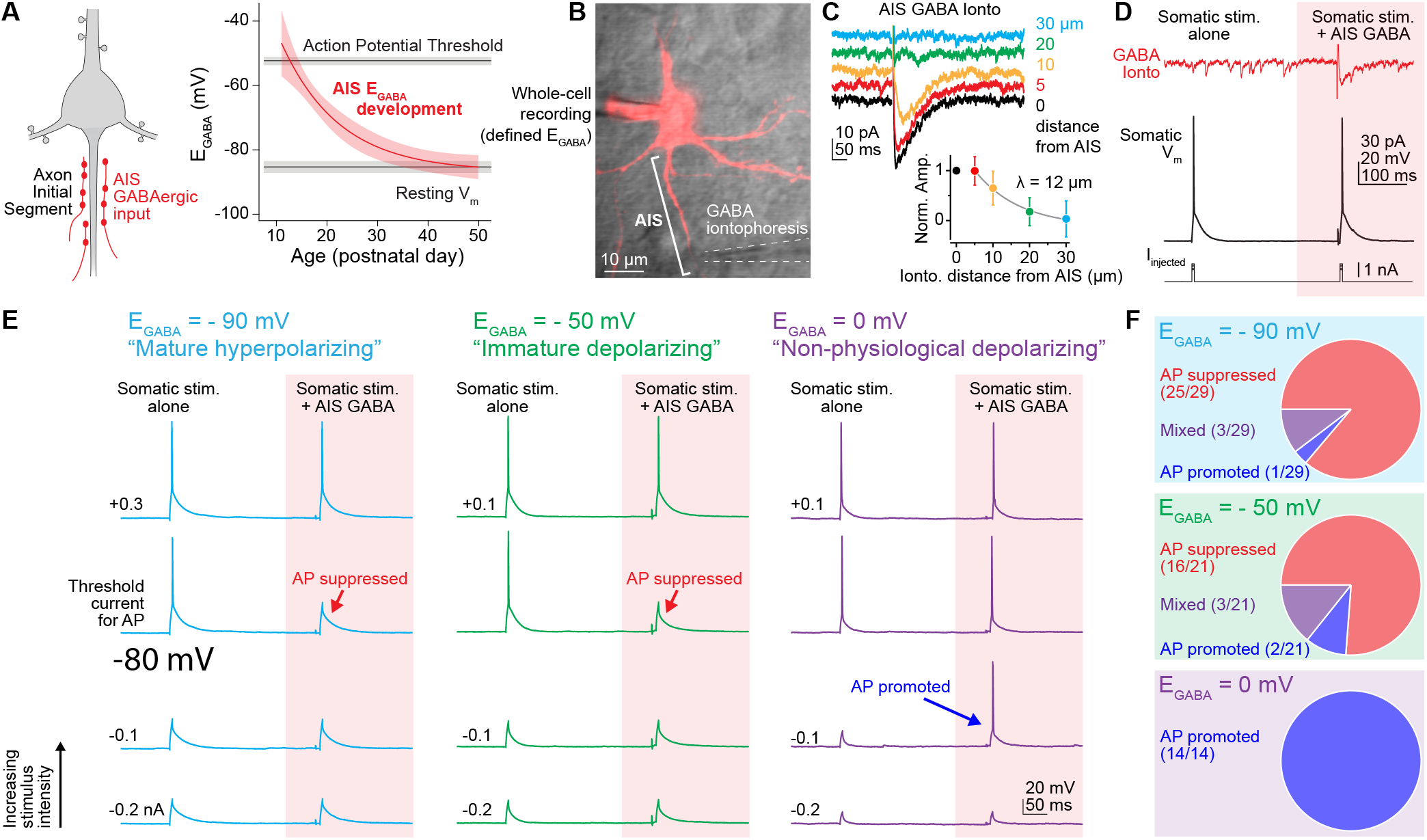
AIS GABA suppresses AP generation at physiological GABA reversal potentials. **A**. Left, chandelier cells form GABAergic synapses on the axon initial segment of layer 2/3 pyramidal neurons in prefrontal cortex. Right, the reversal potential of GABA (E_GABA_) at these synapses follows a periadolescent developmental trajectory, beginning with a depolarized EGABA in early adolescence and reaching a mature, hyperpolarized E_GABA_ by adulthood (P55+). **B**. 2PLSM z-stack of a layer 2/3 pyramidal neuron visualized with Alexa Fluor 594. AIS indicated by bracket. An iontophoresis pipette filled with 1M GABA is placed next to the AIS, near the typical site of GABA synapses. **C**. Spatial spread of GABA after iontophoresis. The amplitudes of GABA PSCs were quantified for different distances of the ionto. pipette from the AIS and fitted with an exponential function. The length constant of GABA spread was 12 μm. **D**. Schematic of experimental protocol. Action potentials (middle) were generated by minimal-stimulation somatic current injection (bottom). To test the effects of GABA_A_ receptor activation on AP generation, GABA iontophoresis was used generate a GABA PSC during the second AP. The relative timing of GABA and the second AP were calibrated so the AP occurred simultaneously with the peak amplitude of the GABA PSC. **E**. GABA iontophoresis suppresses AP generation at physiological values of E_GABA_. Somatic stimulation intensity was calibrated to elicit an action potential in both the presence and absence of GABA (top). When stim. intensity was reduced by 0.1 nA, APs occurring with GABA ionto. were suppressed when E_GABA_ was set to -90 mV (left) or -50 mV (center). In the case of E_GABA_ set to 0 mV, reducing stim. intensity resulted in a suppression of the non-GABA AP but a promotion of the GABA AP (right). With a further decrease in stim. intensity, all APs were suppressed across conditions (bottom). **F**. The proportion of cells that showed AP suppression, AP promotion, or a combination of both patterns with GABA ionto. across different E_GABA_ values.

Multiple chandelier cells synapse on each pyramidal cell AIS, with each chandelier cell forming multiple synapses. Chandelier cells are thought to fire in bursts that are coordinated across the chandelier cell network (Inan and Anderson, 2014; Schneider-Mizell et al., 2021; Viney et al., 2013). Current transgenic approaches do not allow for complete labeling of this population without leak to genetically-similar basket cells that synapse onto non-AIS targets(Schneider-Mizell et al., 2021; Taniguchi et al., 2013). To study how concerted recruitment of all GABA_A_ synapses on the AIS affects AP initiation, we mimicked signaling from multiple chandelier cells with GABA iontophoresis. GABA release was spatiotemporally limited by using an iontophoresis amplifier with pipette capacitance compensation circuitry. GABA spread was limited to ∼20 μm from the pipette tip (space constant: 12 ± 2.85 μm, n = 8, **Fig. 1*C***).

To compare AP generation with and without GABA, two APs were evoked by minimal-stimulation somatic current injection (0.6 – 2.5 nA, 5 ms duration, 400–500 ms inter-AP interval). GABAergic currents (PSCs) were evoked via iontophoresis to coincide with the second AP (**Fig. 1*D***). To ensure the maximum effect of GABA application, the relative timing of GABA and the second AP were calibrated so the AP occurred when the the presynaptic cell and focus on the postsynaptic effects of altering E_GABA_, GABA PSC peaked in amplitude. With an E_GABA_set to −90 mV, as seen in the mature AIS, GABA suppressed AP generation. Surprisingly, when E_GABA_ set to -50 mV, as seen in the immature AIS, GABA continued to suppress AP generation (**Fig. 1*E-F***). To test if AIS GABA could drive AP generation under more overt depolarizing conditions, we set E_GABA_ to 0 mV, a condition that leads to a non-physiologically strong driving force on Cl_-_. In these conditions case, APs were promoted by pairing current injection with AIS GABA application (**Fig. 1*E-F***). This confirms that GABA can act in an excitatory fashion with regards to AP initiation, albeit at E_GABA_ values far more depolarized than those observed over periadolescent development.

### Physiologically relevant E_GABA_ at the AIS both delay AP initiation and increase AP threshold

While GABA applied to the AIS consistently suppresses APs initiation across physiological E_GABA_ values, the mechanism by which this AP failure occurs may differ across conditions. Given the importance of the AIS in setting the voltage threshold for AP generation (Kole et al., 2008), we hypothesized that AIS GABA decreases AP probability by increasing AP threshold, which would likely also delay AP timing. To determine if AIS GABA application is associated with changes in AP threshold or timing, we compared APs generated in the presence and absence of GABA iontophoresis in the same three E_GABA_ conditions as in Fig. 1. For pairs of action potentials, we measured AP threshold and onset timing with and without GABA application and examined differences on a trial-by-trial basis to control for subtle changes in recording conditions.

For GABA iontophoresis onto the initial segment, setting E_GABA_ to −90 mV led to a delay in AP initiation. Without GABA, AP onset occurred 3.64 ms (IQR: 3.46 − 4.21) after the onset of somatic current injection. By contrast, AIS GABA delayed AP onset to 4.28 ms (IQR 4.18 − 4.48) after current injection, a within-sweep median difference of 0.52 ms (IQR: 0.280.85) (**Fig. 2*A, D***). AP onset was similarly delayed when E _GABA_was set to −50 mV. APs without GABA application occurred at 3.68 ms (IQR: 3.49 – 4.08) after current injection and were delayed to 4.15 ms (IQR: 3.674 – 4.3) with GABA (within-sweep median difference = 0.26 ms, IQR: 0.03 – 0.40, **Fig. 2*A, D***). By contrast, non-physiological 0 mV E _GABA_was associated with an advancement of AP onset, with APs occurring 0.85 ms (IQR: -1.23 – -0.48) earlier with AIS GABA (**Fig. 2*A, D***).

**Figure 2:**
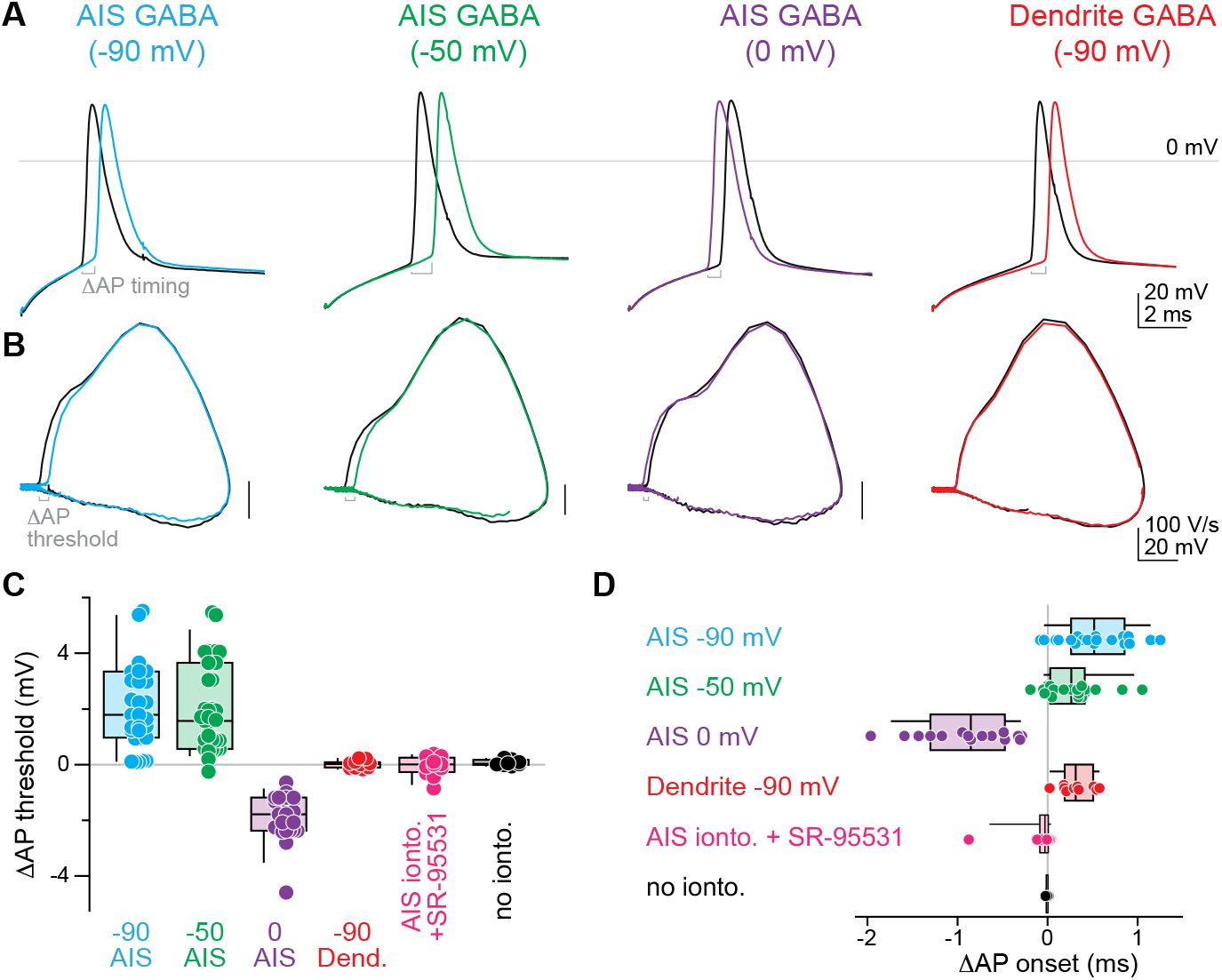
GABA alters AP threshold and AP onset differentially across synaptic location. **A**. AP onset is altered by GABA application to the AIS or dendrite. Black: APs with no GABA input, color: APs with GABA. **B**. Phase plane plots (dV/dt vs Vm) of APs with and without GABA reveal a location-specific effect of GABA on AP threshold. **C**. Summary of the effects of GABA on AP threshold. **D**. Summary of the effects of GABA on AP onset.

To determine whether these effects were due to a specific role for AIS GABAergic synapses, we next compared these results to similar experiments targeting a dendritic branch proximal to the soma. These proximal dendrites are targets of basket cells (Taniguchi et al., 2013), In contrast to GABAergic synapses onto the AIS, dendritic GABAergic synapses exhibit mature E_GABA_ values by P12. Therefore, we set E_GABA_ to -90 mV for all experiments with dendritic GABA application. With dendritic GABA, the median within-sweep change in AP onset was 0.32 ms (IQR: 0.17 – 0.46), with control APs reaching threshold at 4.23 ms (IQR: 3.75 – 4.61) after current injection and dendritic GABA AP onset at 4.63 ms (IQR: 4.11 – 4.76) (**Fig. 2*A, D***). This is consistent with dendritic GABA_A_ receptors acting as a sink for somatic current injections, delaying AP onset.

Next, we examined changes in AP threshold with and without GABA application at the AIS and dendrite for all E_GABA_ values. For GABA application at the AIS when E_GABA_ was set to −90 mV, median AP threshold across cells shifted from −59.4 mV (IQR: −61.6 – −56.9) without GABA to −57.9 mV (IQR: −59.9 – -54.1) with GABA (**Fig. 2B**). Within-sweep changes in AP threshold had a median value of 1.79 mV (IQR: 1.09 – 3.18) (**Fig. 2C**). When E_GABA_ was set to −50 mV, AIS GABA application also resulted in an increase in AP threshold, from −54.5 mV (IQR: −56.7 – −52.6) without GABA to −52.1 mV (IQR: −54.9 – −50.7) with GABA (**Fig. 2*B***). The median within-sweep change in threshold with E_GABA_ = −50 mV was 1.57 mV (IQR: 0.57 – 3.35) (**Fig. 2*C***). 0 mV E_GABA_ again showed a divergent effect, hyperpolarizing AP threshold by 1.78 mV (IQR: −2.36 – -1.23, **Fig. 2*B, C***). By contrast, dendritic GABA application did not result in a change in AP threshold (baseline threshold = −56.6 mV, IQR: −57.8 – -55.4; GABA threshold = −56.9 mV, IQR: −57.9 – -55.2, within-sweep median = 0.04 mV, IQR: 0.01 –0.08) (**Fig. 2*B, C***). To confirm that all effects were due to GABA signaling itself, and not simply changes in intrinsic properties that occurred between the two APs, we examined the pair of APs without applying GABA or with GABA_A_ receptors blocked with SR95531 (10 μM). Neither AP threshold nor timing were altered in these cases (without iontophoresis: median change in AP threshold = 0.03 mV, IQR: −0.001 – 0.11; median change in AP onset = -0.005 ms, IQR: −0.009 -90 mV led to a delay in AP initiation. Without GABA, AP onset occurred 3.64 ms (IQR: 3.46 – 4.21) after the onset of somatic current injection. By contrast, AIS GABA delayed AP onset to 4.28 ms (IQR 4.18 – 4.48) after current injection, a within-sweep median difference of 0.52 ms (IQR: 0.28 −0.002502; SR95531: median change in AP threshold = 0 mV, IQR: -0.18 – 0.15, median change in AP onset = −0.02 ms, IQR: −0.07 – 0.001). Overall, these data indicate that initial segment GABA has an inhibitory effect on AP initiation across E_GABA_ values observed in periadolescent development. Additionally, the stability AP threshold when GABA was applied to proximal dendrites highlights the privileged control of AIS GABA over AP threshold.

### Ca _V_3 and K_V_7 channels do not play a role in AP threshold changes during AIS GABA application

We next sought to test whether specific ion channels may play a role in GABA-mediated AP threshold changes. K_V_7 channels are present at the AIS and contribute to setting the resting membrane potential and threshold in hippocampal granule cells and prefrontal pyramidal neurons (Battefeld et al., 2014; Hu and Bean, 2018; Martinello et al., 2015). Cholinergic input can shift AP threshold in hippocampal granule cells via Ca-dependent inhibition of axonal K_V_7 (Martinello et al., 2015). To determine if K_V_7 channel activity affects AP threshold relative to GABA application, we applied the K_V_7 antagonist XE991 (10 μM) during the same protocol as described above. GABA continued to alter AP threshold in both cases during XE991 application (E_GABA_ = −90 mV, median change in AP threshold = 3.63 mV, IQR: 1.86 – 5.57; E_GABA_ = −50 mV, median change in AP threshold = 2.05 mV, IQR: 0.70 – 2.83), suggesting that K_V_7 channels do not contribute to threshold changes after brief AIS GABA stimulation in these neurons (**Fig. 3*A***). Similarly, we saw no alteration in changes in AP onset during XE991 application (E_GABA_ = −90 mV, median change in AP onset = 0.84 ms, IQR: 0.50 – 1.22; E_GABA_= -50 mV, median change in AP onset = 0.44 ms, IQR: 0.27 – 1.15, **Fig. 3*B***).

**Figure 3:**
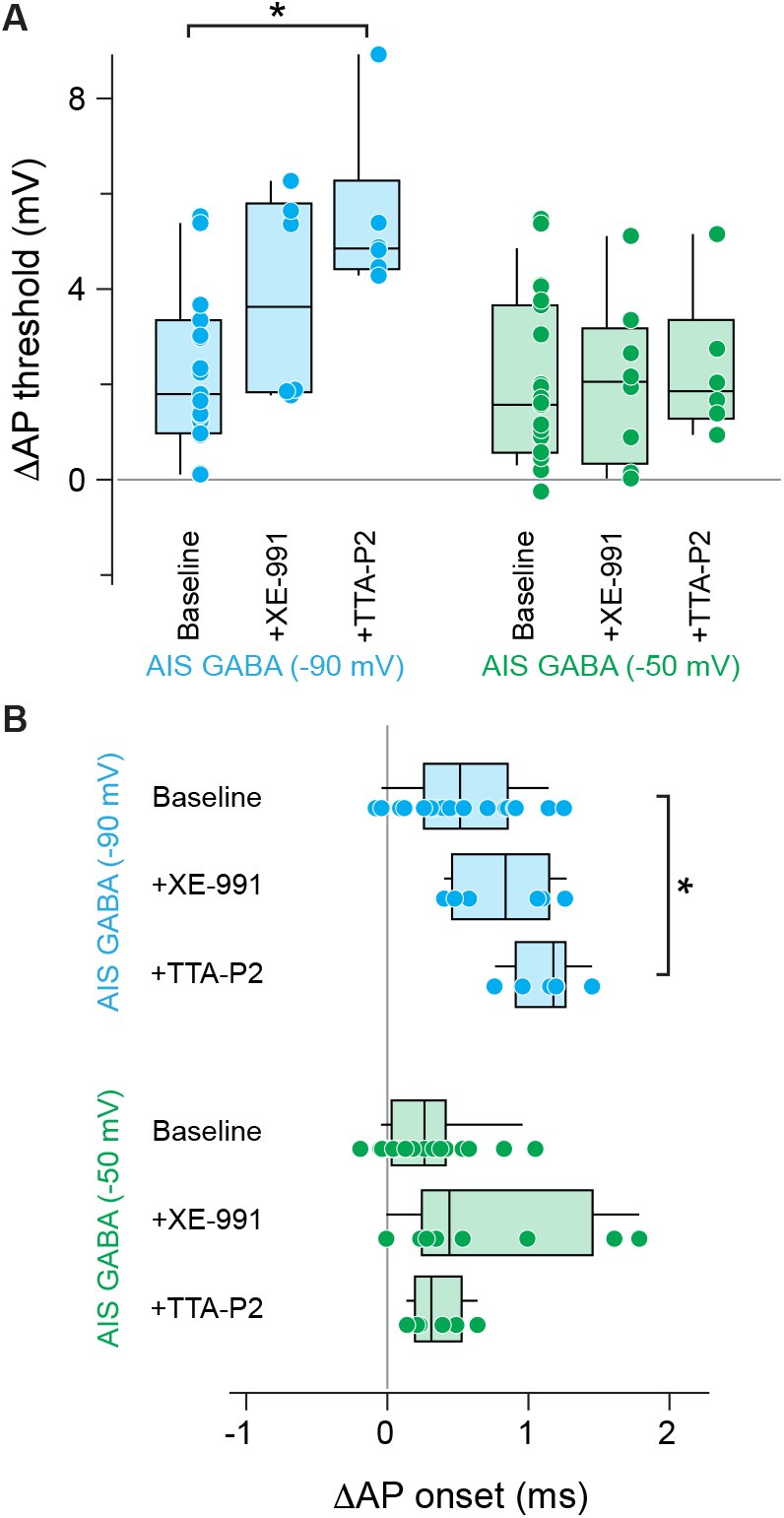
AIS GABA effects on threshold and timing are independent of Ca_V_3 and K_V_7 channels. A. Summary of the effects of GABA on AP threshold in the presence of different ion channel antagonists. XE-991 was applied at 10 μM concentration; TTA-P2 was applied at 2 μM. ^*^ p <0.05 Wilcoxon Rank Sum Test. B. Summary of the effects of GABA on AP onset in the presence of different ion channel antagonists. ^*^ p <0.05 Wilcoxon Rank Sum Test.

Beyond Na_V_ and K_V_ channels, the AIS is also populated by calcium channels (Ca_V_s). Low-voltage activated Ca_V_3 channels are found at the AIS in a variety of cell classes (Bender and Trussell, 2009; Bender et al., 2012; Clarkson et al., 2017; Dumenieu et al., 2018; Fukaya et al., 2018; Hu and Bean, 2018; Lipkin et al., 2021; Martinello et al., 2015). These channels activate and inactivate near resting membrane potential and can be relieved from inactivation by membrane hyperpolarization (Huguenard, 1996). Therefore, small changes in membrane voltage can lead to large shifts in the number of Ca_V_3 that can open during subsequent depolarization. To assess if Ca_V_3 channel availability contributes to the observed changes in AP threshold and onset, we blocked Ca_V_3 channels with the selective antagonist TTA-P2 (2 μM). As with XE991, TTA-P2 did not block AIS GABA-induced changes in AP threshold (E_GABA_ = −90 mV, median change in AP threshold = 4.85 mV, IQR: 4.55 – 5.26; E_GABA_ = −50 mV, median change in AP threshold = 1.85 mV, IQR: 1.46 – 2.57) or AP onset (E_GABA_ = −90 mV, median change in AP onset = 1.18 ms, IQR: 1.00 – 1.20; E_GABA_ = −50 mV, median change in AP onset = 0.31 ms, IQR: 0.22 – 0.47) in either E_GABA_ condition (**Fig. 3*A, B***). In fact, TTA-P2 application with hyperpolarized E_GABA_ led to a small but significant increase in AP threshold and delay in AP initiation (threshold: p = 0.001, onset: p = 0.0019, Mann-Whitney Wilcoxon Test). This is consistent with the observed role of AIS Ca_V_3 channels in affecting AP threshold (Bender and Trussell, 2009). In the absence of GABA iontophoresis, AP threshold was hyperpolarized from −54.4 mV (IQR: −56 – −54.1) to -59.4 mV (IQR: -61.6 – -56.9) by TTA-P2 application alone. Having ruled out the dependence of AIS GABA-mediated AP threshold changes on Ca_V_3 and K_V_7, we next focused on the role of shunting inhibition in the observed effects.

### AIS GABA modulates AP threshold via shunting inhibition across physiological E_GABA_ values

GABAergic inhibition of pyramidal cell firing can occur through two major mechanisms: hyperpolarizing inhibition and shunting inhibition. Inhibition that is hyperpolarizing shifts local membrane potential from its current state to values even further from AP threshold. By contrast, shunting inhibition does not alter membrane potential, but does lower local membrane resistivity, making it more difficult to change membrane voltage away from E_GABA_. Shunting inhibition can occur across a range of E_GABA_ values and resting membrane potentials, but shunting lasts only as long GABA_A_ receptors are open (Paulus and Rothwell, 2016). Overt changes in membrane potential may accompany shunting inhibition if E_GABA_ is far from the baseline membrane potential. These changes can outlast the shunt, as they depend on the membrane time constant. As our pharmacological approaches ruled out several ion channels recruited by changes in membrane potential in GABA-mediated threshold and timing shifts, we sought to understand the role of shunting inhibition in these effects instead.

If shunting inhibition underlies changes in AP threshold, these changes would be predicted to remain at a similar magnitude regardless of the recorded neuron’s holding potential. Conversely, changes in threshold that rely on GABA-mediated changes in membrane potential will increase with larger driving force on GABA and decrease with smaller driving force (**Fig. 4*B***). To test GABA effects at different holding potentials, we added an additional pair of APs that were generated from a more depolarized resting membrane potential. As before, the second AP of each set occurred during the peak GABA current (**Fig. 4*A***). When E_GABA_ was set to −50 mV, the driving force (DF) for GABA was lower when the cell was held at −65 mV compared to -85 mV (a decrease in GABA PSP amplitude is visible in **Fig. 4*A***). Despite this decrease in driving force, however, changes in AP threshold were consistent across holding potentials (high DF: median change in AP threshold = 1.57 mV, IQR: 0.57 – 3.35; low DF: median change in AP threshold = 1.53 mV, IQR: 0.69 – 2.41, **Fig. 4*C-F***), suggesting that shunting inhibition is a major contributor to AIS GABA-driven changes in AP threshold. A similar pattern was seen when E_GABA_ was -90 mV. Despite the increased driving force on GABA at −65 mV compared to -85 mV, we observed no increase in the AIS GABA-mediated AP threshold change withE_GABA_at −90 mV (low DF: median change in AP threshold = 1.79 mV, IQR: 1.09 – 3.18; high DF: median change in AP threshold = 1.82 mV, IQR: 1.48 – 3.10, **Fig. 4*C-E, G***). Overall, as changes in AP threshold were consistent across different driving forces, these changes following GABAergic input to the AIS appear to largely result from shunting inhibition.

**Figure 4:**
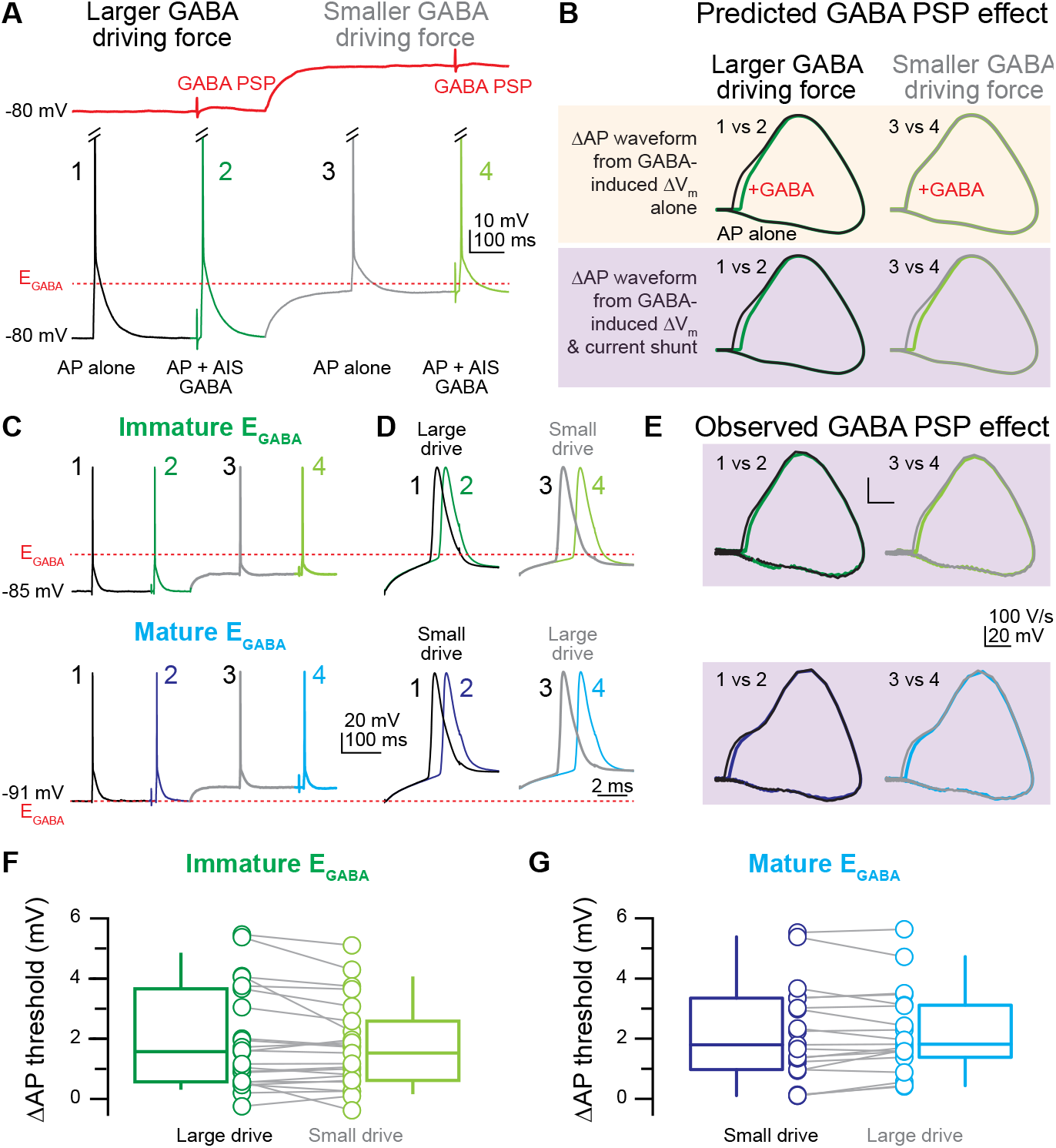
GABA depolarizes AP threshold across E_GABA_ values via shunting inhibition. **A**.Schematic of experimental protocol. GABA PSCs were elicited by GABA iontophoresis onto the AIS (top) at two different membrane potentials to alter the driving force on GABA according to E_GABA_ (dashed red line). Pairs of action potentials were evoked by somatic current injection at each membrane potential. GABA iontophoresis was delivered during the second AP of each pair. AP threshold was compared between APs generated with and without GABA from the same holding current (i.e. AP 2 (dark green) was compared to AP 1 (black)). **B**. Predicted effects of GABA PSP driving force on AP threshold. If GABA-mediated changes in membrane potential underlie changes in AP threshold, reducing the driving force by depolarizing the baseline Vm would be predicted to reduce the change in threshold (yellow, top). If GABA-mediated shunting inhibition underlies changes in AP threshold, the change in threshold would be predicted to stay the same across high and low driving forces (purple, bottom).**C**. Top, example APs elicited from a representative neuron with “immature” E_GABA_. Bottom, example APs elicited from a representative neuron with “mature” E_GABA_. E_GABA_ for each neuron is represented as a dashed red line. **D**. Expanded time base shows AP waveform of spikes elicited with and without AIS GABA at different GABA driving forces. Top, E_GABA_ = −50 mV, bottom, E_GABA_ = −90 mV. **E**. Phase plane plots of APs displayed in C and D. Black/gray, without GABA; green/blue, with GABA. **F**. Summary of the effects of AIS GABA on AP threshold across different GABA driving forces when E_GABA_ was set to −50 mV. Dark green, high driving force; light green, low driving force. **G**. Summary of the effects of AIS GABA on AP threshold across different GABA driving forces when E_GABA_ was set to −90 mV. Dark blue, low driving force; light blue, high driving force.

### Reduced Na_V_ availability from depolarizing AIS GABA contributes to GABA-mediated threshold changes

Shunting inhibition appears to play a large role in AIS GABA-mediated changes in AP threshold. However, it remains possible that membrane depolarization in the “immature” E_GABA_condition may additionally contribute to changes in AP threshold. For example, the fraction of AIS Na_V_s that are “available” for AP electrogenesis—the fraction of channels that are currently closed rather than in the inactivated or open state—is voltage dependent and changes markedly at voltages just subthreshold for AP initiation. Thus, small changes in membrane potential can translate to large changes in Na_V_ availability and therefore AP threshold. However, exploring the role of Na_V_ pharmacologically is difficult, as interfering with Na_V_ function necessarily interferes with AP generation. One elegant method for exploring the role of Na_V_ function following depolarizing GABAergic input is to decouple the shunting and depolarizing effects of GABA on AP waveform. Previous work performed in axonal bleb recordings from the substantia nigra demonstrated that depolarizing GABA inputs lead to slower APs via a reduction in the number sodium channels in the closed state (Kramer et al., 2020). The authors determined that this effect was independent of GABA-mediated shunting by evoking depolarization with direct current injection rather than GABA_A_ receptor activation. While GABA_A_ activation was more effective at altering AP height than current injection alone, the comparable effectiveness of GABA application and current injection on reducing AP speed suggested that GABA-mediated depolarization rather than shunting was the main driver of the observed effect on AP speed.

We used this approach to quantify the relative contributions of these two actions of depolarizing GABA on AP threshold in our layer 2/3 pyramidal neurons. Using the same approach as in Fig. 4, we correlated the effects of membrane depolarization with changes in AP threshold. First, we quantified the difference in membrane potential before each evoked AP (**Fig. 5*A***, center). We then measured the difference in AP threshold between the first APs in each pair, which were generated without GABA but from different baseline potentials (**Fig. 5*A***, right). To compare GABA-mediated changes to those generated by membrane depolarization alone, we also quantified the amplitude of the recorded GABA PSP in both physiological E_GABA_ conditions. This allowed us to plot the ΔV_m_ against the ΔAP threshold in each condition: current injection alone, “immature” E_GABA_, and “mature” E_GABA_. We predicted that pure shunting effects would fall along the Y-axis of the resulting plot, as changes in threshold would occur in the total absence of V_m_ changes. Conversely, we predicted that changes in threshold that resulted from reduced Na_V_ availability at depolarized V_m_ would be positively correlated and fall in the upper righthand quadrant (**Fig. 5*B***).

**Figure 5:**
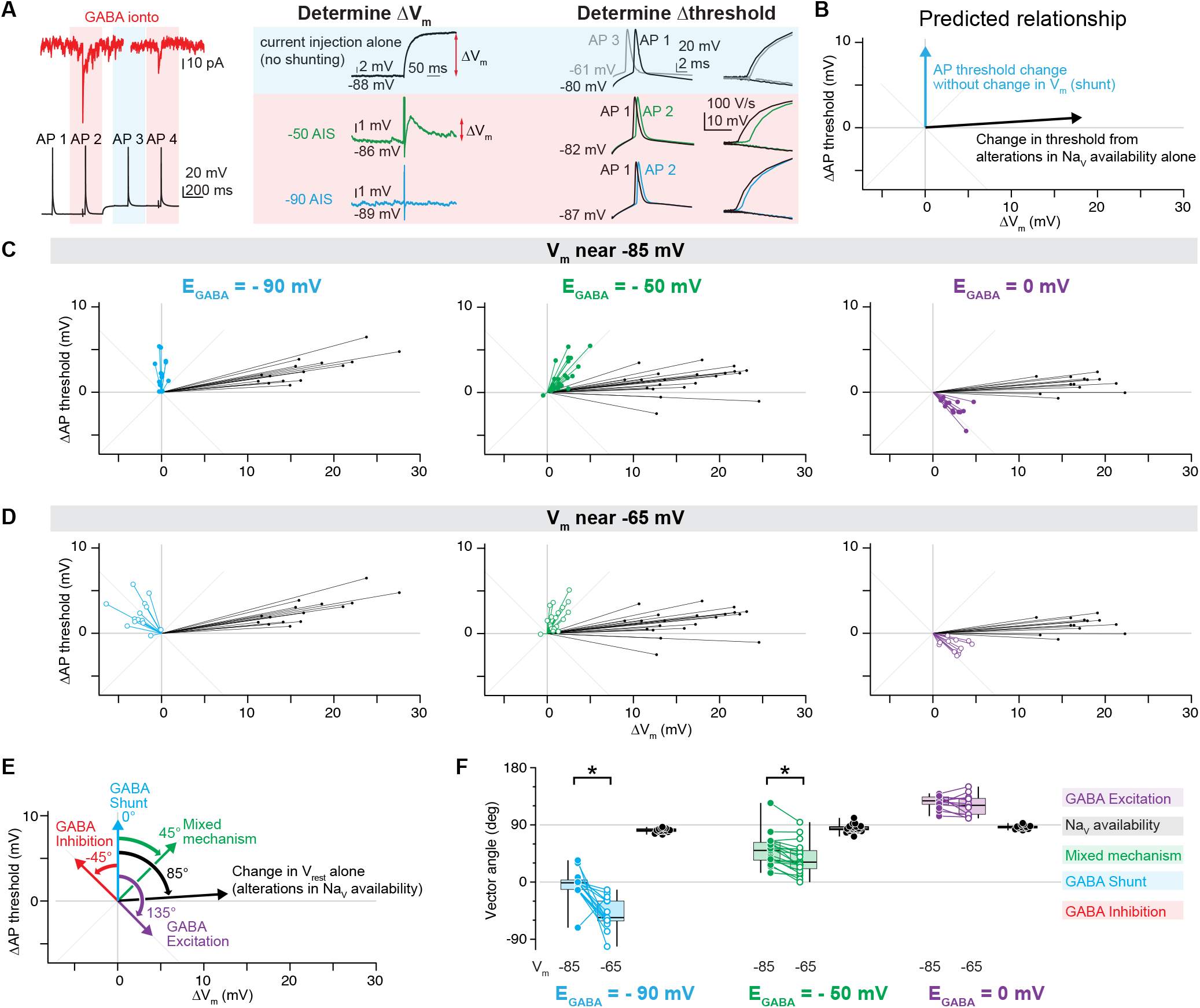
Sodium channel availability contributes to GABA-mediated threshold changes. **A**. Schematic of experimental protocol used to determine the relationship between changes in V_m_ and changes in AP threshold with and without shunting inhibition. Top left, example GABA postsynaptic currents (red) recorded from -88 mV (left) and -68 mV (right) Bottom left, APs generated from different membrane potentials. Center, depiction of how changes in Vm were calculated for each condition. Note that no change in Vm occurred with -90 mV AIS _EGABA._ Right, depiction of how changes in AP threshold were calculated for each condition. For current injection alone, AP1 and AP3 (both without GABA) were compared. **B**. Predicted relationship between changes in Vm and changes in AP threshold. **C**. For neurons held at -85 mV holding potential, changes in AP threshold plotted against changes in Vm for each condition. **D**. For neurons held at -65 mV holding potential, changes in AP threshold plotted against changes in Vm for each condition. **E**. The relationship between ΔVm and Δthreshold can be quantified by measuring the angle of each point from the y-axis. An 85° angle described NaV availability changes due to membrane depolarization alone. An angle of 0° is indicative of a pure GABA shunt. Angles near -45° were considered GABA inhibition. Angles near 135° demonstrated a purely excitatory effect of GABA. Finally, an angle of 45° suggested a mixed mechanism, with some contributions of shunting and some reduction in NaV availability. **F**. For each point in C and D, vector angle from the y-axis was calculated and plotted by condition and by baseline Vm. ^*^ p <0.05 Wilcoxon Rank Sum Test.

When E_GABA_ was set to -90 mV and GABA was applied to the AIS, the GABA-associated change in V_m_ was minimal, as V_m_ (-85 mV) was very close to E_GABA_ and DF was therefore low. However, as we observed a pronounced depolarization of AP threshold, plotting Δthreshold against ΔV_m_ resulted in points falling along the Y-axis, in line with our predictions of shunting inhibition as the primary driver of AP threshold depolarization in “mature” E_GABA_ conditions (**Fig. 5*C***). In contrast to the purely shunting effect of “mature” E_GABA_, plotting Δthreshold against ΔV_m_ in the “immature” E_GABA_ condition revealed a mixed effect of shunting and reductions in Na_V_ availability, as data points clustered between the vertical axis (a pure shunt) and the line describing Na_V_ availability changes (**Fig. 5*C***). In the case of non-physiological E_GABA_, where GABAergic input to the AIS was capable of promoting AP generation through hyperpolarization of AP threshold (**Fig. 1-2**), changes in V_m_ were negatively correlated to changes in AP threshold and the resultant points were found in the bottom right cluster of the graph (**Fig. 5*C***). Patterns were largely similar when APs were evoked from a more depolarized membrane potential (-65 mV), which differentially altered driving force on GABA in different E_GABA_ conditions (**Fig. 5*D***). However, rather than falling along the vertical axis, “mature” E_GABA_ data points now were plotted in the upper lefthand axis, indicative of hyperpolarization-mediated inhibition (**Fig. 5*D***).

To determine if differences existed between E_GABA_ conditions at each baseline V_m_, we quantified the above results by measuring the angle of each plotted point to the Y-axis (**Fig. 5*E***). We found that the vector angle differed significantly between holding potentials for “mature” E_GABA_ (p = 0.0015, Wilcoxon Ranked-Sum Test), as well as between holding potentials for “immature” E_GABA_ (p = 0.0006, Wilcoxon Ranked-Sum Test, **Fig. 5*F***). In the “immature” E_GABA_ condition, the angle was smaller when driving force was reduced, indicating a stronger, but not completely explanatory, effect of shunting inhibition on the observed changes in AP threshold. Taken as a whole, these data indicate that GABA synapses on the AIS of pyramidal neurons inhibit AP generation across the developmental range of E_GABA_ by depolarizing AP threshold. In immature pyramidal neurons, where E_GABA_ is depolarized relative to resting membrane potential, this effect is achieved through a combination of shunting inhibition and depolarization-mediated reductions in sodium channel availability. By adulthood, when E_GABA_ reaches its mature, hyperpolarized value, shunting inhibition is the dominant mode by which axo-axonic synapses regulate AP initiation.

## DISCUSSION

Here, we show the mechanism by which GABAergic input onto the axon initial segment regulates AP initiation. While E_GABA_ undergoes a developmental switch from depolarized to hyperpolarized in L2/3 neuron initial segments (Rinetti-Vargas et al., 2017), the overall effect of GABA at the AIS is inhibitory across periadolescent E_GABA_ ranges. This uniformly inhibitory effect of GABA on AP initiation is largely mediated by shunting inhibition, where changes in membrane conductance associated with the opening of GABA_A_ receptors reduce the efficacy of incoming currents to generate APs. The effect of this shunt is a depolarization of AP threshold when GABA is applied to the AIS, as well as a delay in AP onset. Dendritically-applied GABA delays AP onset with no effect on AP threshold. Together, these effects suggest that AIS GABA, presumably from chandelier cells, is preferentially poised to inhibit AP generation.

### Depolarized axonal E_GABA_ leads to inhibitory effects on AP initiation

The axon initial segment is a boundary region in neurons that separates the somatodendritic region from the axonal region while harboring characteristics of both (Bender and Trussell, 2012; Leterrier, 2018; Nakada et al., 2003). Dendritic E_GABA_ is almost exclusively hyperpolarized relative to the cell’s resting membrane potential, while axonal E_GABA_, particularly at axonal terminals, is depolarized relative to rest (Price and Trussell, 2006; Ruiz et al., 2010; Turecek and Trussell, 2001; Zorrilla de San Martin et al., 2017). The AIS shows a mixed phenotype, with E_GABA_ depolarized through early adolescence and subsequently switching to hyperpolarized in adulthood (Rinetti-Vargas et al., 2017). GABA’s reversal potential is set by the activity of two chloride transporters, NKCC1 and KCC2, which import and export chloride respectively (Blaesse et al., 2009). It has been hypothesized that KCC2 is excluded from the axonal domain, resulting in depolarized E_GABA_ throughout the axon (Báldi et al., 2010; Hedstrom et al., 2008; Szabadics et al., 2006). However, the AIS likely has KCC2 activity in adulthood, as evidenced by its periadolescent switch (Rinetti-Vargas et al., 2017). Thus, the AIS has some properties of both somatodendritic and axonal regions.

Understanding how E_GABA_ is regulated is a necessary component of GABAergic synapse function at the AIS; however, GABA’s inhibitory or excitatory effects are not purely governed by E_GABA_ alone. Work exploring the effect of GABA_A_ receptors expressed further down the axon highlights how difficult it is to predict the effects of axonal GABA from E_GABA_ alone. In cerebellar Purkinje neuron terminals and hippocampal mossy fiber boutons, E_GABA_ is depolarized relative to resting membrane potential and leads to increases in neurotransmitter release via increases in AP-associated bouton calcium influx (Ruiz et al., 2010; Zorrilla de San Martin et al., 2017). At the presynaptic Calyx of Held, however, a depolarized E_Cl_ does not alter the amplitude of the postsynaptic response (Price and Trussell, 2006). In the axons of highly-branched midbrain dopamine neurons, GABA’s reversal potential is elevated relative to rest, but GABA application or agonism leads to a shortening and slowing of APs due to a combination of depolarization-mediated sodium channel inactivation and shunting inhibition (Kramer et al., 2020). In L2/3 pyramidal neurons of immature rat somatosensory cortex, paired recordings between chandelier and pyramidal cells found that chandelier inputs did not promote AP generation in most cases; however, a small fraction of chandelier cell APs did directly generate APs in their postsynaptic targets when E_GABA_ was set to -48 mV in pyramidal cell whole-cell recordings (Szabadics et al., 2006). In this case, the ability to generate APs seemed related to GABA PSC amplitude, with larger events correlated with AP promotion. However, for a similar range of GABA PSC amplitudes, we did not observe a similar pattern of direct AP generation in prefrontal cortex. While E_GABA_ is depolarized in each of these examples, the responses to axonal GABA vary across cell types and synaptic location. Understanding how the geometry of the axon as well as the conductances present in the bouton or along the axon influence AP waveform and transmitter release is crucial to predicting the postsynaptic effects of GABA in a particular cell type.

Membrane potential changes following GABA_A_ receptor activation may alter the availability of ion channels near the site of GABA input. In the AIS, Na_V_s, low-threshold Ca_V_s, and K_V_s all show some availability at resting membrane potential, suggesting that small, GABA-mediated changes in membrane potential may lead to the opening or closing of these channels that may promote excitation or inhibition. We show that in immature pyramidal initial segments, GABA-mediated depolarization reduces the availability of sodium channels, working in concert with shunting to reduce AP threshold (**Fig. 5**). Additionally, our data suggest that K_V_ 7 channels and Ca_V_ 3 channels are not involved in changing AP threshold after GABA input to the AIS in this manner (**Fig. 3**). However, this does not rule out the possibility that longer-lasting changes in postsynaptic membrane potential or calcium signaling may occur downstream of these channels. For example, hyperpolarization-activated cyclic nucleotide-gated (HCN) channels, which underlie I_h_ current, have been shown to contribute to excitatory effects on AP generation following hyperpolarizing IPSPs in CA1 pyramidal neurons (Kwag and Paulsen, 2009). Specifically, IPSPs advanced AP onset by approximately 10 ms when the IPSP occurred on the falling phase of the theta oscillation. While threshold changes were not examined directly in that study, this example shows that GABA may have effects across a range of time courses. As I_h_ does not appear present in L2/3 pyramidal neurons (Larkum et al., 2007), we did not explore the contribution of HCN channels here. Our results here provide mechanistic insight into one sliver of that range: the effects on AP threshold and onset. However, it does not preclude the possibility that hyperpolarized and depolarized E_GABA_ have different downstream effects that we have not captured here. Modeling data comparing shunting and hyperpolarizing GABA in a network show that while hyperpolarizing inhibition is more effective at synchronizing firing across neurons, shunting inhibition increases the network’s ability to distinguish between different stimuli (Burman et al., 2023). Additionally, work revealing an adolescent switch in the homeostatic plasticity of chandelier cell synapses suggests that opposing downstream effects of depolarizing and hyperpolarizing GABA must play a role in governing plasticity at this synapse (Pan-Vazquez et al., 2020).

### Depolarized E_GABA_ and AIS homeostatic plasticity

In a recent paper, Pan-Vasquez, Wefelmeyer, et al. observe a switch in the direction of homeostatic plasticity at chandelier cell synapses in L2/3 of somatosensory cortex (Pan-Vazquez et al., 2020): when the authors increased network activity in this brain region before the GABAergic switch (P12–P18), they observed a decrease in the number of chandelier cell synapses, while this same manipulation resulted in an increase in chandelier cell synapses during adulthood (P40–P46). Thus, they were able to characterize homeostatic plasticity rules across this developmental switch and showed that different values of E_GABA_ do have different downstream effects. Therefore, it is surprising to see that regardless of E_GABA_ value (within the physiological range, at least), the effect of AIS GABA on threshold is uniformly inhibitory, driving threshold to more depolarized potentials. These differences may result from differences in brain regions between prefrontal and somatosensory cortices, or that ongoing synaptic activity *in vivo* may have additional effects on E_GABA_ that are not evident in quiescent *ex vivo* slice conditions (Burman et al., 2023). Additionally, it may be that changes in AP threshold are not the main driver of homeostatic plasticity at this synapse, and may instead depend on, for example, slower changes in calcium influx at the AIS (Lipkin and Bender, 2020), or the soma (Evans et al., 2013; Grubb and Burrone, 2010) that are affected by subthreshold AIS voltage. In line with this, homeostatic, activity-dependent plasticity of AIS position persists after blockade of APs but disappears with block of calcium channels (Grubb and Burrone, 2010). As more research emerges showing the role of E_GABA_ shifts in governing behaviors from sleep pressure to migraine-related pain (Alfonsa et al., 2022; Paige et al., 2022), the precise consequences of E_GABA_ changes across a range of excitability and plasticity metrics will need to be explored.

### Dendritic and axonal GABA provide differential control over AP generation

Basket cells and chandelier cells are two morphologically-distinct classes of parvalbumin-positive interneurons that synapse onto the perisomatic region of pyramidal cells in the prefrontal cortex (Taniguchi et al., 2013). Basket cells provide inhibitory input onto the soma and proximal dendrites, whereas chandelier cells specifically target the axon initial segment (Taniguchi et al., 2013). While parvalbumin-positive interneurons broadly are crucial in orchestrating gamma oscillations (Cardin et al., 2009; Sohal et al., 2009), the individual roles of basket and chandelier cells in this oscillatory activity have been difficult to disentangle. Here, we show that chandelier cell synapses on the AIS have preferential control over AP threshold, while both dendritic and AIS GABA synapses can alter AP onset. A similar phenotype was observed in dentate granule cells, where somatic application of the GABA_A_ agonist muscimol resulted in reductions in AP peak amplitude and delays in AP onset without affecting AP threshold (Rojas et al., 2011). In the same cells, muscimol application to the AIS depolarized AP threshold across E_GABA_ values. These differential effects on AP generation translated to alterations in EPSP summation depending on GABA_A_ receptor location: somatic GABA receptors created a large membrane shunt, which reduced the efficacy of each EPSP in driving the neuron towards threshold, while initial segment GABA receptors raised AP threshold without altering the efficacy of each EPSP. In the dentate gyrus, this effectively raises the number of coincident inputs required to convey information into the hippocampus (Rojas et al., 2011). Further experiments are needed to explore how basket cell and chandelier cell synapses may shape temporal summation in pyramidal cells in the same manner in the prefrontal cortex.

## METHODS

### Electrophysiology and Imaging

All procedures were performed in accordance with University of California Animal Care and Use Committee guidelines. C57Bl/6 mice of either sex, aged postnatal day 24–50 were used for all experiments. Mice were anesthetized and acute coronal slices (250 μm) containing medial prefrontal cortex were collected. Cutting solution contained: 2.5 mM KCl, 1.2 mM NaH2PO4, 30 mM NaHCO3, 20 mM HEPES, 5 mM Na+ ascorbate, 2 mM thiourea, 3 mM Na+ pyruvate, 25 mM glucose, 10 mM MgSO4 • 7 H20, 0.5 mM CaCl2 • 2 H20, and 92 mM NaCl (pH 7.3, 300 mOsm). Slices were then allowed to recover for 15 min at 32-34° C in an N-methyl D-glucamine (NMDG) recovery solution containing 2.5 mM KCl, 1.2 mM NaH2PO4, 30 mM NaHCO3, 20 mM HEPES, 5 mM Na+ ascorbate, 2 mM thiourea, 3 mM Na+ pyruvate, 25 mM glucose, 10 mM MgSO4 • 7 H20, 0.5 mM CaCl2 • 2 H20, and 93 mM NMDG (pH 7.3, 300 mOsm). Following this recovery, slices were allowed to equilibrate in HEPES cutting solution at room temperature for 2 hours before recording. All solutions were bubbled with 5% CO2/95% O2.

Whole-cell recordings were made from layer 2/3 pyramidal neurons in prefrontal cortex using Dodt contrast optics. Patch electrodes were pulled from Schott 8250 glass (3-4 MΩ tip resistance) and filled with internal solution. Three different internal solutions were used to set the GABA reversal potential (EGABA) for each experiment. For EGABA = -90 mV experiments, internal solution contained: 113 mM K-gluconate, 9 mM HEPES, 2.25 mM MgCl2, 2.25 mM MgSO4, 10 mM sucrose, 14 mM Tris2-phosphocreatine, 4 mM Na2-ATP, 0.3 Tris-GTP, and 0.1 μM EGTA. For EGABA = -50 mV experiments, internal solution contained: 102 mM K-gluconate, 11 mM KCl, 9 mM HEPES, 4.5 mM MgCl2, 10 mM sucrose, 14 mM Tris2-phosphocreatine, 4 mM Na2-ATP, 0.3 Tris-GTP, and 0.1 μM EGTA. For EGABA = 0 mV experiments, internal solution contained: 113 mM KCl, 9 mM HEPES, 4.5 mM MgCl2, 10 mM sucrose, 14 mM Tris2-phosphocreatine, 4 mM Na2-ATP, 0.3 Tris-GTP and 0.1 μM EGTA. For all internal solutions, pH was 7.2-7.5 and osmolarity was adjusted to ∼290 mOsm. To visualize the AIS with two-photon imaging, all internals were supplemented with 20 μM Alexa-594. After whole-cell configuration was achieved, cells were allowed to equilibrate for at least 5 minutes to allow for whole-cell dialysis of the internal solution.

Electrophysiological data were acquired using a Multiclamp 700B amplifier (Molecular Devices) and custom routines written in IgorPro (Wavemetrics). Data were acquired at 50 kHz and filtered at 20 kHz. APs were generated with somatic current injection and injection current amplitude was calibrated to be just-suprathreshold. Briefly, somatic current step amplitude was adjusted in 100 pA increments until the minimum value where APs were reliably generated could be found. AP threshold was defined as the voltage at which dV/dt exceeded 15 mV/ms. AP onset was defined as the time at which threshold was reached after the start of the current injection. Changes in threshold and onset were averaged over 5-50 sweeps for each cell. To minimize electrode drift within the slice, all recordings were made using a quartz electrode holder (Sutter Instrument). Cells were excluded if series resistance exceeded 20 MΩ. Data were corrected post hoc for junction potential as follows: EGABA = -90 mV, 12 mV; EGABA = -50 mV, 8 mV; EGABA = 0 mV, −5 mV.

GABA iontophoresis was performed using parafilm-wrapped borosilicate pipettes (0.5 μm tips, 1M GABA in H2O [pH 4]) placed in proximity to neurites visualized with two-photon laser-scanning microscopy. Two-photon laser-scanning microscopy was supported by either a Coherent Ultra II laser or Mira 900 laser was tuned to 800 nm, and Alexa-594 was visualized with R9110 photomultiplier tubes positioned to capture epifluorescence (<660 nm light, no bandpass) and transfluorescence (535/150 bandpass, Semrock) through a 40x, 0.8 NA objective paired with a 1.4 NA oil immersion condenser (Olympus). After placement near the neurite of interest, GABA was released onto the neurite using an iontophoresis amplifier with pipette capacitance compensation circuitry (MVCS-C, npi electronic, Germany) (30-400 nA application current, -40 nA retention current, 2 ms pulses). EGABA was confirmed for each cell in voltage-clamp. Experiments were performed in the presence of CGP55845 (10 μM) to block GABAB receptors.

## Chemicals

SR95531, CGP55845, and XE991 were from Tocris Bioscience. TTA-P2 was from Alomone Labs. Alexa Fluor 594 was from Invitrogen. All other chemicals were from Sigma.

### Statistics

Unless otherwise specified, all data are reported as medians with interquartile ranges (IQR) in text and displayed with box plots showing the median, quartiles, and 10%/90% tails). n denotes number of cells. Sample sizes were chose based on standards in the field. No assumptions were made about data distributions, and unless otherwise noted, two-side, rank-based nonparametric tests were used. Significance level was set for an α value of 0.05. Holm-Sidak correction was used for multiple comparisons when appropriate. Statistical analysis was performed using IgorPro 8.0 (Wavemetrics).

## Acknowledgments

We are grateful to members of the Bender and Nelson labs at UCSF and to Drs. Z Khaliq and P. Kramer for insightful comments and discussion. This work was supported by National Science Foundation Grants 1650113 (to A.M.L.) and National Institutes of Health Grants AA027023 and MH112729 (to K.J.B.).

## Author contributions

A.M.L. and K.J.B. designed research; A.M.L. performed research; A.M.L. analyzed data; A.M.L. and K.J.B. wrote the paper.

